# Predicting an Individual’s Cerebellar Activity from Functional Connectivity Fingerprints

**DOI:** 10.1101/2023.03.18.533265

**Authors:** Vaibhav Tripathi, David C Somers

## Abstract

The cerebellum is gaining scientific attention as a key neural substrate of cognitive function; however, individual differences in the cerebellar organization have not yet been well studied. Individual differences in functional brain organization can be closely tied to individual differences in brain connectivity. ‘Connectome Fingerprinting’ is a modeling approach that predicts an individual’s brain activity from their connectome. Here, we extend ‘Connectome Fingerprinting’ (CF) to the cerebellum. We examined functional MRI data from 160 subjects (98 females) of the Human Connectome Project young adult dataset. For each of seven cognitive task paradigms, we constructed CF models from task activation maps and resting-state cortico-cerebellar functional connectomes, using a set of training subjects. For each model, we then predicted task activation in novel individual subjects, using their resting-state functional connectomes. In each cognitive paradigm, the CF models predicted individual subject cerebellar activity patterns with significantly greater precision than did predictions from the group average task activation. Examination of the CF models revealed that the cortico-cerebellar connections that carried the most information were those made with the non-motor portions of the cerebral cortex. These results demonstrate that the fine-scale functional connectivity between the cerebral cortex and cerebellum carries important information about individual differences in cerebellar functional organization. Additionally, CF modeling may be useful in the examination of patients with cerebellar dysfunction, since model predictions require only resting-state fMRI data which is more easily obtained than task fMRI.

## Introduction

Although traditionally viewed primarily as a motor structure, the cerebellum supports a broad range of non-motor cognitive functions, including working memory, attention, language and higher cognition (Stoodley & Schmahmann, 2009; Schmahmann, 2019; Brissenden et al., 2021). This view is supported both by studies of patients with cerebellar damage and by neuroimaging studies of healthy subjects (Schmahmann et al., 2007; Stoodley et al., 2012). Brain networks supporting specific forms of cognition comprise not only cerebral cortical regions, but also cerebellar regions and there’s growing evidence for fine-scale cerebro-cerebellar connectivity (Buckner et al., 2011; Liu et al., 2022). The fact that cerebro-cerebellar anatomical connectivity is not monosynaptic complicates the analysis of these networks. Descending connections pass via the pons and ascending connections pass via thalamus (Steriade & Llinas, 1988). Resting-state functional MRI has proven an effective way to reveal multisynaptic brain networks generally (Gordon et al., 2017) and cerebro-cerebellar networks more specifically (Guell et al., 2018).

Most neuroimaging investigations of cerebellar function have been focused on group-level analyses. In contrast, individual subject-level analysis offers a number of potential advantages, including the ability to observe fine-scale structures that would be blurred by group analyses and the ability to develop precision medicine diagnostics for individual patients (Braga & Buckner, 2017; Somers et al., 2021; Xue et al., 2021). Here we extend individual subject research methods that have been applied to the cerebral cortex to the cerebellum.

Passingham and colleagues proposed that each cortical area has a unique pattern of cortico-cortical connections – a *‘connectional fingerprint’* – that could be used to functionally localize cortical areas in individuals (Mars et al., 2018). Connectome Fingerprinting (CF) is a computational neuroimaging technique that combines non-invasive connectome measurements to predict fine-scale functional brain organization in individual subjects (Osher et al., 2016; Saygin et al., 2012; Tobyne et al., 2018). CF modeling approaches based on connectomes derived from structural or resting-state functional MRI have been successfully applied throughout the cerebral cortex; however, the cerebellum has received little attention to date.

Connectome Fingerprinting predicts voxel-wise activations to a particular task from the resting state functional connectivity from voxels in a search space to regions in a parcellation scheme. Utilizing high quality resting state data, we can predict with sufficient accuracy the activations in out of sample subjects (Tavor et al., 2016) even with low n datasets (Osher et al., 2019; Tobyne et al., 2018).

We utilized task and resting-state fMRI data from the Human Connectome Project (HCP) young adult dataset (n=160). The HCP examined seven cognitive paradigms that probed a diverse set of brain networks: Working memory, Gambling, Motor, Language, Social, Relational, Emotion. Each task compared an experimental condition with a control condition as well as a fixation baseline condition. The Working Memory and Motor tasks also included sub-conditions, with different categories of visual images in the working memory task and different body parts in the motor task. Group level task activation in these paradigms is reported in Barch et al. (2013). We found that activations predicted by CF models at an individual level outperformed the group level task activation as a predictor for out of sample subjects. The subject’s own functional connectivity predicted task activation better than others suggesting individual specificity of the model and the coefficients of the model were similar to the task activations in the cortex which were not a part of the model suggesting a close link between cerebellar-cortical connectivity and cortical functional activations.

## Methods

### Dataset

We used 160 subjects (98 females) from the Human Connectome Project Young Adult (HCP-YA) dataset (Van Essen et al., 2012) who had acquisitions on both 3T and 7T scanners. The HCP Young Adult initiative collected high quality structural, resting state and task data on a population of young adults (ages 22-35 years). The study was approved by the Washington University Institutional Review Board and all subjects gave informed consent for the study. The dataset is available to all on HCP’s data management platform, ConnectomeDB (https://db.humanconnectome.org). A custom Siemens CONNECTOM Skyra MRI Scanner was used. All subjects participated in two days of scanning which included a high-resolution structural T1 weighted MRI, T2 weighted MRI. During resting-state scans (32 channel head coil, voxel resolution = 2 mm isotropic, in-plane FOV = 208 × 180 mm, 72 slices, multi-band factor 8, TR = 720 ms, TE = 33.1 ms, 1200 TRs) subjects were asked to visually fixate on a cross and do nothing, in particular. Resting-state scans consisted of four runs of fifteen minutes each collected in two separate sessions. The task runs had the same acquisition protocol but differed in the number of frames (TRs). Half of the task and resting-state runs were acquired using left-to-right phase encoding and another half on the right-to-left phase encoding.

The subjects participated in seven task experiments in the scanner that examined different aspects of cognition: Working memory, Gambling, Motor, Language, Social, Relational, Emotion. The first six tasks consistently yielded robust activation of portions of the cerebellum. Only a very small portion of the cerebellum was activated by the Emotion Task, so we consider this task only in the supplemental materials. We describe the tasks briefly here, details can be found in the initial HCP task fMRI paper (Barch et al., 2013). The working memory task involved a ‘2-back’ working memory condition in which subjects were asked to report when the current stimulus matched a stimulus presented two trials prior, and a ‘0-back’ control condition in which subjects were asked to report when a presented stimulus matched a target that was presented at the start of the block. Stimuli were presented in blocks and across blocks four different categories of images were employed: places, faces, body parts and tools. There were 8 task blocks per run, half of the blocks were 2-back, and the other half were 0-back. Task blocks were 25 s long with 10 trials per block. There were also 4 fixation blocks per run of 15 s each. Total 405 TRs were collected per run. We explored the 2-back vs 0-back contrast, and the 2-back body, 2-back face, 2-back place, and 2-back tool conditions each contrasted with their 0-back counterpart.

The Gambling task compared reward processing to loss processing using a shared task paradigm; the key manipulation was to covertly change the odds of rewards/losses across blocks of trials. On each trial, a mystery card was presented on the screen and the subject had to guess if the number on the card would be higher or lower than five.

Reward blocks had mostly reward trials (6 out of 8, and others could be loss and/or neutral trials) whereas Punishment blocks had mostly loss trials (6 out of 8, and others could be neutral and/or reward trials). Each run had two blocks of reward and punishment 28 s each and four fixation blocks of 15 s each and a total of 253 TRs. We explored the Punishment-Fixation and Reward-Fixation conditions.

The Motor task involved visual cues where subjects were asked to tap either their left or right fingers, squeeze either their left or right toes or move the tongue. The blocks were 12 s long and each run consisted of two blocks for each movement (Left Hand, Right Hand, Left Foot, Right Foot, and Tongue) and three fixation blocks of 15 s each. Total of 284 TRs per run were collected. We analyzed each body part vs average (AVG) contrasts for all five conditions (LH, LH, RF, RH, T) where the average included the other 4 conditions. The Language task consisted of four ‘story’ task and four ‘math’ task blocks per run. The ‘story’ condition blocks consisted of stories from Aesop’s fables and subjects were asked to respond via button press to a 2-alternative forced-choice (2-AFC) question about the topic of the story. The ‘math’ condition blocks consisted of auditory presentations of math calculations like “eight plus five equals” followed by two possible answers; subjects reported the correct answer via button press. The duration of the blocks were fixed at 30 s, some subjects performed the math computations faster, so they were given additional trials to match the duration of the story blocks. Each run was 316 TRs long and we studied the Math vs Story contrast for the language task.

The Social Cognition Task or ‘Theory of Mind’ task incorporated video clips of 20 s long of simple shapes (e.g., squares, circles and triangles) that moved across a background. In the ‘Theory of Mind’ condition, the coordinated movement of the shapes suggested a social interaction between them, while in the control condition the shapes moved about independently of one another. Each run had five video blocks, a total of ten across the two runs (five for each of the two conditions). The subjects were asked to report whether they observed a social interaction, no interaction or were not sure about the interaction. Each run had 5 fixation blocks of 15 s each and a total of 274 TRs and we examined the Theory of Mind vs Fixation and Random Movement vs Fixation contrasts.

The Relational Task asked subjects to infer a relationship between two objects and to examine that relationship between another pair of objects. Stimuli consisted of shapes that had a certain texture. In the relational task condition, subjects had to compare two shapes on the top of the screen with each other to determine whether they differed in shape or texture and then report whether the two shapes on the bottom also differed along the same dimension or not. In the matching control condition, subjects were presented with two shapes on the top, one on the bottom and a cue (shape or texture) and were asked to report whether the bottom shape was similar to either of the top shapes in the cued dimension. Each run had three relational and three match blocks that lasted 18 s and three fixation blocks of 16 s length, with a total of 232 TRs and we analyzed the Relational Task vs fixation and the Matching Task vs fixation contrasts.

The Emotion task asked subjects to examine facial expressions of emotions. Stimuli consisted of faces (experimental) and shapes (control) conditions during which subjects were presented with two faces/shapes on the top and one face/shape on the bottom and they were ask to match the bottom face/shape with the top ones and report via button press which top face/shape was similar to the bottom one. The faces had either angry or fearful expressions. Each run consisted of three faces and three shapes blocks (all 18 s long) and had a total of 176 TRs and we looked over the Faces vs fixation and Shapes vs fixation contrasts.

### Preprocessing

We used the minimally preprocessed data as available on the HCP database portal. The preprocessing pipeline (Glasser et al., 2013) included corrections for artifact, gradient non-linearity correction, motion and EPI distortion followed by temporal denoising and bandpass filtering (0.001 - 0.008 Hz). The structural and functional images were registered from the native space to MNI space. Freesurfer pipeline (Dale et al., 1999; Fischl et al., 1999) was used to convert it to the 168k fsaverage space which was converted to the 32k CIFTI “grayordinates” space which included the two cerebral hemispheres as surfaces and the subcortical regions as volumes registered in the MNI 2mm space. The data was spatially smoothed with a 2 mm FWHM. We demeaned the resting state data across time within each voxel/vertex and regressed out the mean global signal and concatenated the four runs resulting in 60 minutes (total 4800 TRs) of resting-state data per subject.

### GLM Analysis

General Linear Model analysis of the task data was performed using modified scripts (https://github.com/Washington-University/HCPpipelines/tree/master/TaskfMRIAnalysis) as made available by the HCP consortium (Barch et al., 2013; Glasser et al., 2016). The already analyzed files were downloaded from the analysis section from the HCP database (db.humanconnectome.org). The modified scripts run the FSL-based GLM analysis (Jenkinson et al., 2012) on each voxel/vertex in the 91k grayordinates space. Block-based analysis was performed using a double gamma HRF as implemented in FSL. The contrasts used for various tasks are Working Memory (2BK_BODY vs. fix, 2BK_FACE vs. fix, 2BK_PLACE vs. fix, 2BK_TOOL vs. fix, 2BK vs. 0BK), Gambling (PUNISH vs. fix, REWARD vs. fix), Motor (LH vs AVG, RH vs. AVG, LF vs. AVG, RF vs. AVG, T vs. AVG), Language (MATH vs. STORY), Social (Theory of Mind/TOM vs. fix, RANDOM vs. fix), Relational (REL vs. fix, MATCH vs. fix), Emotion (FACES vs. fix, SHAPES vs. fix).

### Connectivity Fingerprinting

We used the connectivity fingerprinting (CF) approach using resting-state functional connectivity data as described in (Tobyne et al., 2018), illustrated in Figure 1. First, we divided our 160 subjects into test and training datasets consisting of 80 subjects in each group. Our analysis defined 400 regions of interest in the cerebral cortex using the Schaefer 400 parcellation atlas (Schaefer et al., 2017). For a selected contrast, we computed the cerebellar search space as the region with the absolute value of the group average activations greater than 1.5 t-statistic value. The search space for each task contrast defines the set of voxels for which the model will generate predictions. Based on the criteria for the selection of search space, few contrasts (Gambling: PUNISH vs REWARD; Language: MATH, STORY; Social: TOM vs RANDOM; Relational MATCH vs REL; Emotion: FACES vs SHAPES) did not have task-relevant voxels and were not included further in the analysis. For each participant, we computed the subject’s Functional Connectome as a matrix of Pearson correlations between the resting state time courses between each cerebellar voxel and each cortical ROI. ROI time series data were averaged across the voxels of the ROI. Thus for a given task, functional connectome matrices had dimensions of the number voxels in the selected search space x 400 cortical ROIs. We appended the connectivity matrices across the training dataset, and we created a composite predictor matrix. The target matrix was composed of the task activations across the search space appended across the subjects in the training dataset. Using a k-fold (10 folds) ridge regression method as implemented in Scikit Learn (Pedregosa et al., 2011), we trained a model separately for each contrast. The k-folds were used to optimize the hyperparameter in the ridge regression. Using the trained model, we applied our model to the test dataset and computed the prediction accuracy as the Pearson correlation between the actual and the predicted activations. We plotted the predicted activations back on the search space for visualization. To compare the performance of the CF model, we used group average (GroupAvg) as the control model. Group averaged t statistic values in the search space were taken as the prediction for all subjects and compared with the actual activations.

**Figure 1:**
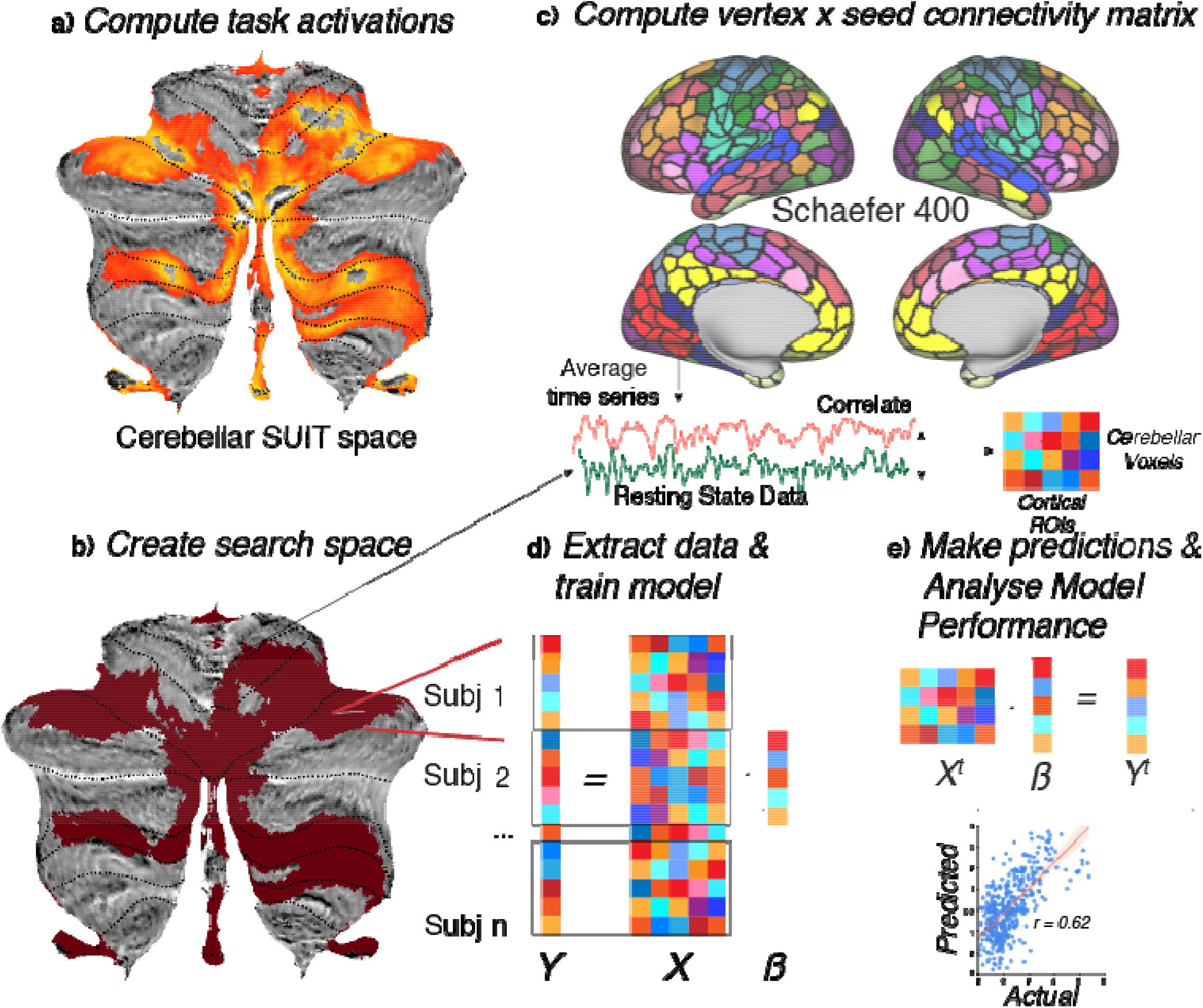
Connectome Fingerprinting Method: We compute the task activations in the cerebellum for a given task and estimate the search space with task-relevant voxels. Then, we create a functional connectivity matrix from the voxels in the search space to cortical parcels defined by a parcellation scheme (here, Schaefer 400 atlas). The connectivity matrices are appended across subjects in the training dataset to create the predictor matrix X. The task activations in the cerebellar search space are appended across training dataset subjects to create the target matrix Y. We then train a ridge regression model with a k-fold cross-validation scheme. The trained model is used to predict activations in the test dataset and the performance is analyzed across the task. The process is repeated for relevant contrasts across the seven tasks of the Human Connectome Project dataset.

We compared the prediction accuracies using the CF and the GroupAvg model using paired t-test as defined in the statsmodels python package (Seabold & Perktold, 2010). Visualizations in the SUIT space (Diedrichsen & Zotow, 2015) were done using the SUITpy python toolbox (https://github.com/diedrichsenlab/SUITPy/releases/latest). To determine if the individual’s functional connectivity was better suited for within-subject prediction, we used the subject’s own functional connectivity (self) and other subjects’ connectivity (others) to predict and compare with the subject’s activation.

The coefficients from the trained model for each contrast were correlated with task activations averaged across the ROIs and subjects to analyze what predictors contributed more to the predictions. The analysis codes were written in Python and used Numpy (Harris et al., 2020), Scipy (Virtanen et al., 2020) and Nipy (https://github.com/nipy/nipy) toolboxes.

## Results

### Prediction accuracy for CF models across various tasks

We analyzed the ability to use resting-state functional connectivity to predict task activations in human cerebellum using the HCP dataset. We first extracted the resting-state time series from each of 400 cerebral cortical ROIs as defined in the Schaefer 400 atlas (Schaefer et al., 2017); each time course was averaged across all voxels within an ROI. The ROI time courses correlated the time series with those of each cerebellar voxel within the task-defined search space. For each task, the cerebellar search space was defined as all cerebellar voxels with robust group-average activations and deactivations, defined as having greater than 1.5 absolute t statistic units. For each subject, the resulting connectivity matrix (voxels x ROIs) constitutes the functional connectome of that subject. A subset of subjects was selected as the ‘training subjects.’

Using the functional connectomes and t-stat task activations across the subjects from the training set (Figure 1), we trained the Connectome Fingerprinting (CF) model using the ridge regression method for each contrast separately (see Methods). We then applied individual subject functional connectomes from subjects in the test set to each model to predict the t-statistics for each voxel in the cerebellar search space for each subject. We evaluated the quality of predictions by examining the correlation between the actual pattern and predicted pattern of t-statistics across the cerebellar search space. We also examined the correlation between each subject’s actual pattern of activation and the group average pattern of activation. This analysis was performed for all subjects of the test set and performance was compared between the CF and group-average predictions.

The Working Memory task visually presented sequences of images and asked subjects to report when a stimulus matched the stimulus shown two stimuli before (2-back task). This was contrasted with a 0-back condition in which subjects were asked to report when a stimulus match a target stimulus presented prior to the block of trials. The 2-back condition created greater working memory demands (e.g., encoding, manipulation) than did the 0-back condition. The CF model predicted individual subject activations in the cerebellum (Figure 2) in the 2-back vs. 0-back (2BK_0BK) contrast significantly better than did the group average approach (CF Mean accuracy=0.30,SD=0.17; GroupAvg M=0.08,SD=0.05; paired t test - t(79) = 12.00, p<0.0001, Cohen’s d=1.78).

**Figure 2:**
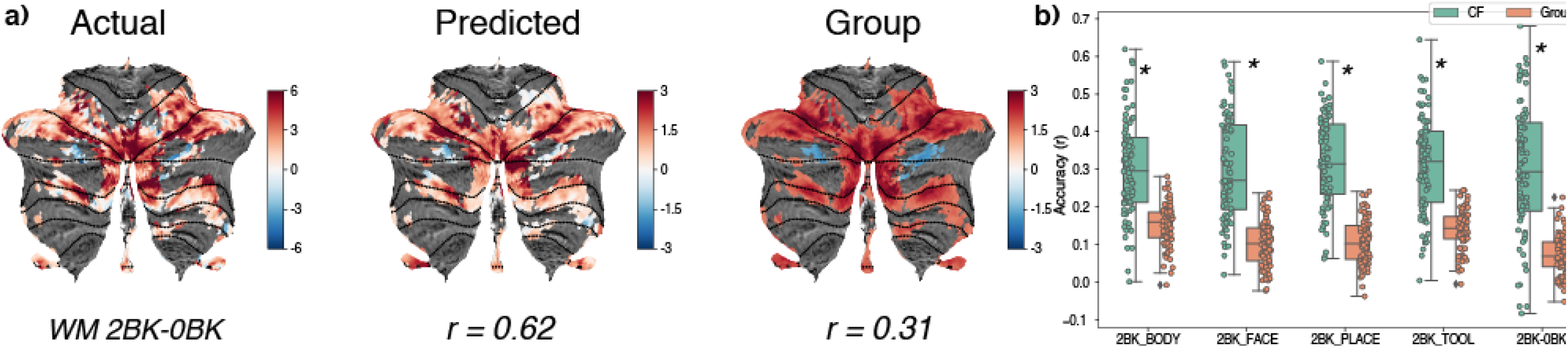
Working memory task predictions. a) Predictions using the CF model (middle panel) and control Group Average model (right panel) for the working memory 2 back vs 0 back contrast. The left panel depicts the actual task activations visualized on the SUIT surface space. b) Prediction accuracy computed as the Pearson correlation coefficient between the predicted and actual activations for the test dataset in the search space for the five contrasts in the working memory task: 2 back Body, 2 back faces, 2 back places, 2 back tools and 2 back vs 0 back. Asterisk depicts statistically significant differences between the prediction accuracies of the two model types.

The 2-back trials were presented in blocks in which the stimuli were restricted to one of four categories of images: bodies, faces, places, or tools. These sub-conditions were each contrasted with the fixation condition, yielding four additional contrasts of interest: 2BK_BODY, 2BK_FACE, 2BK_PLACE, 2BK_TOOL. For each of these contrasts the CF models outperformed the group average predictions: 2BK_BODY (CF M=0.30, SD=0.13; GroupAvg M=0.15, SD=0.06; t(79) = 10.44, p<0.0001, Cohen’s d=1.52); 2BK_FACE (CF M=0.30, SD=0.13; GroupAvg M=0.10,SD=0.06; t(79) = 12.90, p<0.0001, Cohen’s d=1.52); 2BK_PLACE (CF M=0.32, SD=0.12; GroupAvg M=0.11,SD=0.06; t(79) = 15.48, p<0.0001, Cohen’s d=2.26), 2BK_TOOL (CFM=0.31,SD=0.13; GroupAvg M=0.14,SD=0.05; t(79) = 12.01, p<0.0001, Cohen’s d=1.73).

The Gambling task involved reward based processing and the CF model made significant predictions (Figure 3) for the PUNISH (CF (M=0.34, SD=0.15), GroupAvg (M=0.18, SD=0.07); t(79) = 9.49, p<0.0001, Cohen’s d=1.41) and REWARD (CF(M=0.36,SD=0.12), GroupAvg(M=0.08,SD=0.06); t(79) = 18.55, p<0.0001, Cohen’s d=2.75) conditions.

**Figure 3:**
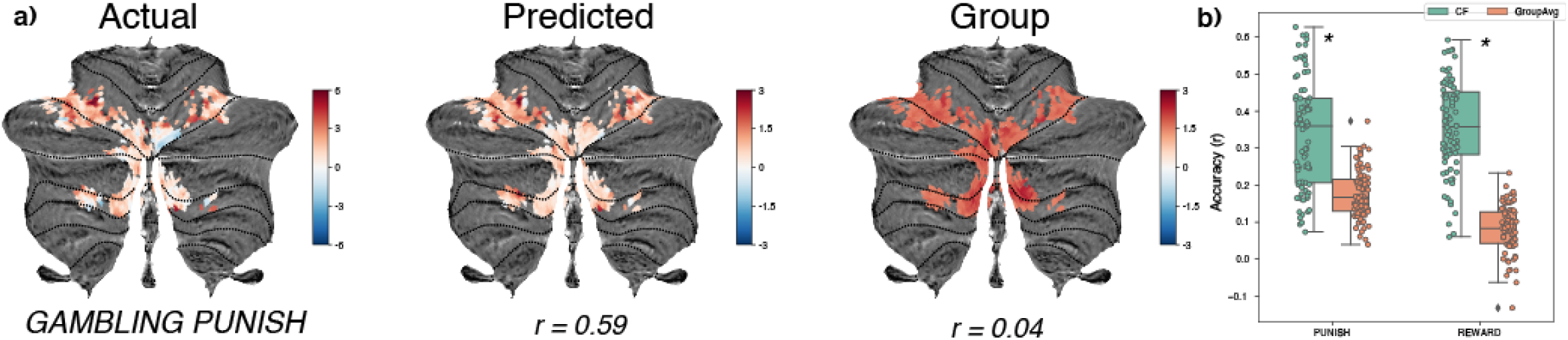
Gambling task predictions. a) Predictions using the CF model (middle panel) and control Group Average model (right panel) for the gambling task punish condition. The left panel depicts the actual task activations. b) Prediction accuracies for the two conditions in the gambling task: Punish (mostly loss trials) and Reward (mostly gain trials). Asterisk depicts statistically significant differences between the prediction accuracies of the two model types.

In the Motor task, subjects had to tap their left hand, left foot, right hand, right foot and tongue during different task blocks. The overall prediction accuracy for the CF models was lower (Figure 4) for the motor contrasts but significant for Left Hand - Average (CF(M=0.16, SD=0.15) GroupAvg (M=0.07, SD=0.11); t(79) = 5.07, p<0.001, Cohen’s d=0.68), RIght Hand - Average (CF(M=0.22, SD=0.17), GroupAvg(M=0.08,SD=0.09); t(79) = 7.41, p<0.0001, Cohen’s d=1.08), Right Foot - Average (CF(M=0.15, SD=0.15), GroupAvg(M=0.05, SD=0.12); t(79) = 4.77, p<0.001, Cohen’s d=0.69) but not significant for Left Foot - Average (CF (M=0.09, SD=0.12), GroupAvg(M=0.07, SD=0.14); t(79) = 1.37, p=0.175, Cohen’s d=0.18) and Tongue - Average (CF(M=0.39,SD=0.15) GroupAvg(M=0.41,SD=0.13); t(79) = -1.66, p=0.100, Cohen’s d=-0.12). In the motor task, the average is computed across all conditions except the contrasted condition, for example, in the Left Hand - Average contrast, the average is computed over the right hand, left and right foot and tongue.

**Figure 4:**
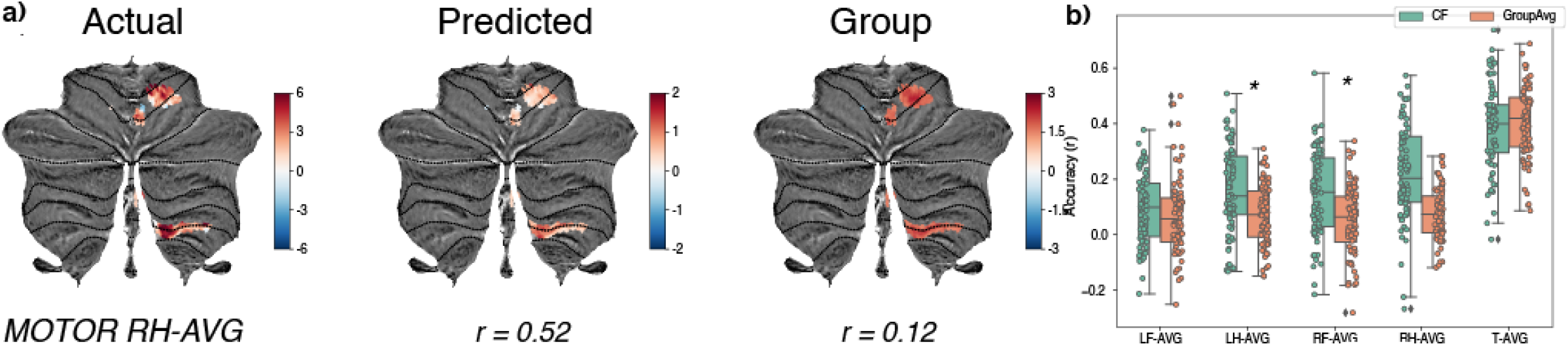
Motor task predictions. a) Predictions using the CF model (middle panel) and control Group Average model (right panel) for the motor task right hand vs average contrast. The left panel depicts the actual task activations. b) In this task subject tapped left and right hand and toes and tongue during a block-based paradigm. Average here refers to average across all except the contrasted one. Prediction accuracies for the five contrasts in the motor task: Left Foot vs Average (LF-AVG), Left Hand vs Average (LH-AVG), Right Foot vs Average (RF-AVG), Right Hand vs Average (RH-AVG) and Tongue vs Average (T-AVG). Asterisk depicts statistically significant differences between the prediction accuracies of the two model types.

In the language task, subjects had to either perform mental calculations in the math condition or comprehend a story and answer true or false questions related to the story. In the MATH-STORY contrast, the prediction accuracy was high for the CF models (FIgure 5) as well as Group average models (CF(M=0.68, SD=0.14), GroupAvg(M=0.68, SD=0.08) but there was no statistical difference between the two models (t(79) = 0.21, p=0.836, Cohen’s d=0.03)

**Figure 5:**
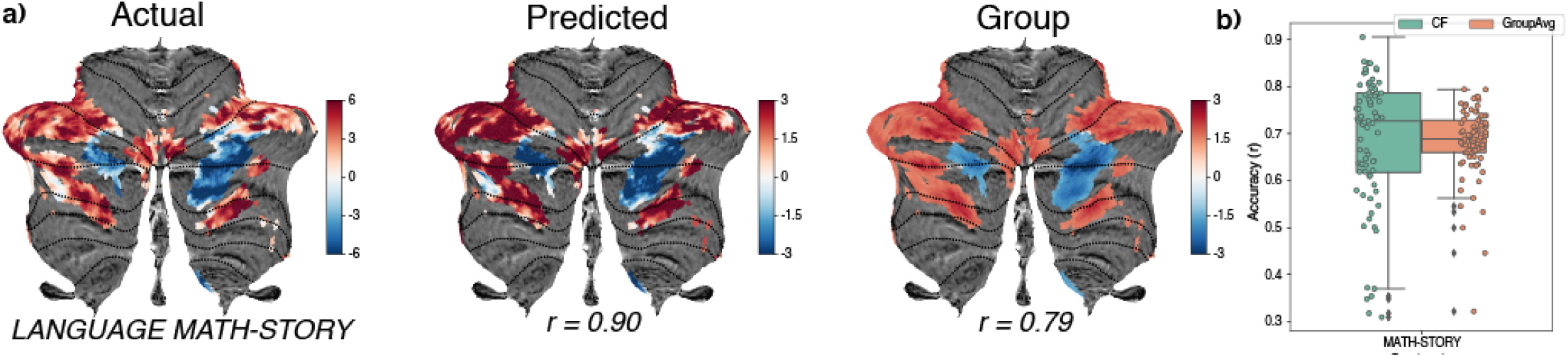
Language task predictions. a) Predictions using the CF model (middle panel) and control Group Average model (right panel) for the language task math vs story contrast. The left panel depicts the actual task activations visualized on the SUIT surface space. b) Prediction accuracy across all subjects using the CF and Group Average control model in the math vs story contrast for the language task where subjects either solve a simple math audio question presented or understand the premise of a short story from Aesop’s fables. Asterisk depicts statistically significant differences between the prediction accuracies of the two model types.

The social task involved looking at the interactions between different shapes moving and making a sense whether the interaction was random or involved the theory of mind (TOM) condition. The prediction accuracy for CF model was significantly higher (Figure 6) as compared to Group Average model for both random (CF(M=0.34, SD=0.12), GroupAvg(M=0.11, SD=0.09); t(79) = 15.56, p<0.0001, Cohen’s d=2.13) and TOM (CF(M=0.36, SD=0.12), GroupAvg(M=0.13, SD=0.08); t(79) = 15.69, p<0.0001, Cohen’s d=2.21) conditions. In the relational task, subjects had to compare shapes or textures of objects presented on the screen whether they related along a dimension or one object match the others along the cued dimension. The CF model (Figure 7) performed significantly better in both Match control (CF(M=0.37, SD=0.11), GroupAvg(M=0.14, SD=0.07); t(79) = 17.82, p<0.0001, Cohen’s d=2.49) and Relational (CF(M=0.40, SD=0.13), GroupAvg(M=0.19, SD=0.06); t(79) = 15.08, p<0.0001, Cohen’s d=2.10) conditions.

**Figure 6:**
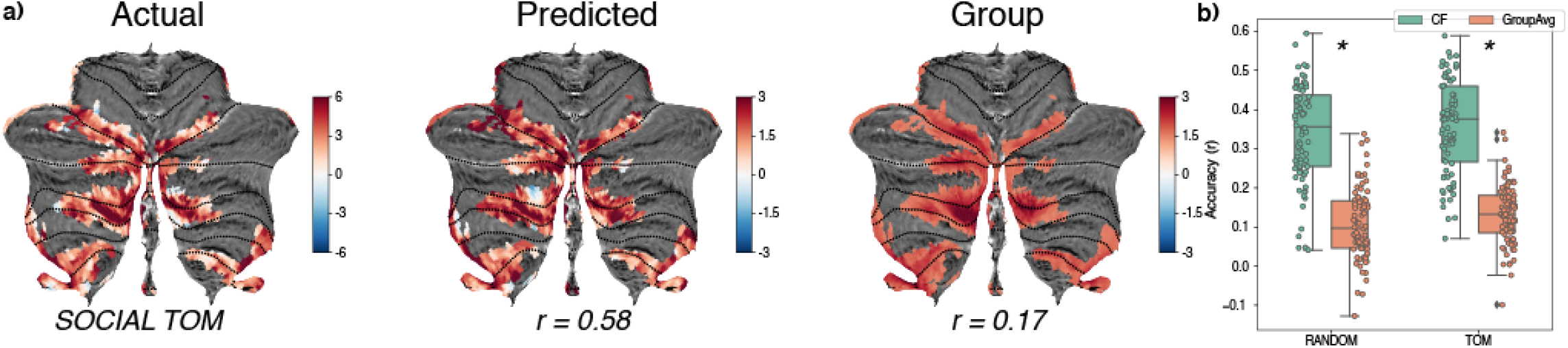
Social task predictions. a) Predictions using the CF model (middle panel) and control Group Average model (right panel) for the social task theory of mind (TOM) condition. The left panel depicts the actual task activations visualized on the SUIT surface space. b) Prediction accuracies across the two model types for all subjects in the random condition where objects on the screen moved randomly or the Theory of Mind (TOM) condition where the objects appeared to have social interaction. Asterisk depicts statistically significant differences between the prediction accuracies of the two model types.

**Figure 7:**
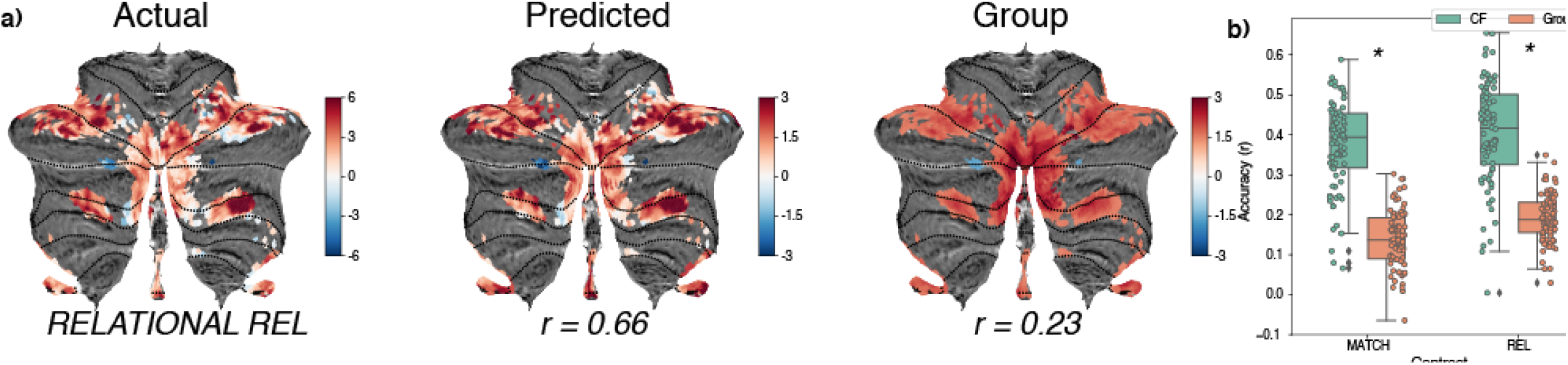
Relational task predictions. a) Predictions using the CF model (middle panel) and control Group Average model (right panel) for the relational task where subjects had to match or compare objects that differed along the dimension of shape or texture. The left panel depicts the actual task activations visualized on the SUIT surface space. b) Prediction accuracies across all subjects in the test dataset compared across the two model types for the match and relational conditions. Asterisk depicts statistically significant differences between the prediction accuracies of the two model types.

The emotion task involved looking at faces with affect and non-affective shapes to elicit and observe the neural responses to emotions. Though the number of task-relevant voxels were less in the cerebellum for the emotion task, CF model (Figure S1) performed significantly better than the Group average model for both the Faces (CF(M=0.35, SD=0.16), GroupAvg(M=0.03, SD=0.14); t(79) = 14.20, p<0.0001, Cohen’s d=2.19) and Shapes conditions (CF(M=0.35, SD=0.34), GroupAvg(M=0.16, SD=0.34); t(79) = 3.48, p<0.001, Cohen’s d=0.54). The Faces-Shapes contrast did not have task-relevant voxels so was not included in the prediction analysis.

### Individual variability in connectivity drives predictions

How does the subjects’ connectivity affect the predictions made through the CF model? We tested the prediction accuracy of the model using different subjects’ resting-state functional connectivity (Figure 8, Figure S2) and compared if it was better than predictions made using subjects’ own functional connectivity which is represented along with the diagonal axis in Fig. 8. We found that for all such pairwise combinations possible for 80 subjects (6320), in more than 98% of combinations the subject’s connectivity made better predictions for the different contrasts in the working memory task (2BK_BODY-98.03 %, 2BK_FACE 98.76 %, 2BK_PLACE 99.14 %, 2BK_TOOL 0.98.37). The 2BK-0BK contrast had a reduced value as 93.24 %. The contrasts in the motor tasks had reduced performance within-subject connectivity vs across (LF-AVG 69.09%, LH-AVG 77.08 %, RF-AVG 76.37 %, RH-AVG 87.92 %, T-AVG 89.81 %). The two conditions in the Gambling task also had a very high degree of within-subject connectivity performance (PUNISH - 97.92 %, REWARD - 98.98 %). We saw similar results with the Social task (RANDOM - 98.65 %, TOM 99.38 %) and the Relational task (MATCH 99.25 %, REL 98.97 %). The Emotion task shapes contrast did around 93.11 %, and Language MATH-STORY 92.86 %. The two-sided Kolmogorov-Smirnov statistic for all contrasts except MOTOR LF-AVG came out greater than 0.35, with all p < 10^−10^.

**Figure 8:**
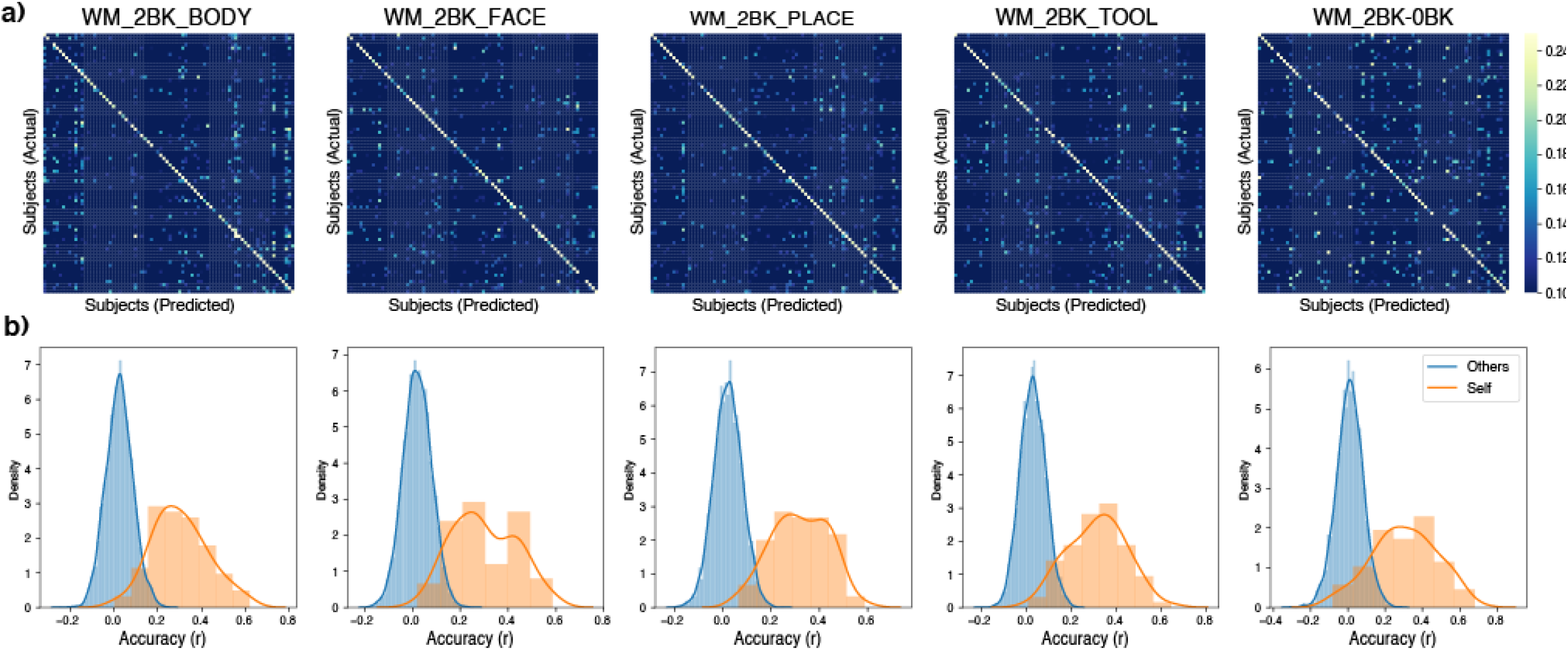
Cross connectivity predictions: a) Predicting a subject’s task activation using their own functional connectivity (self) or other subjects’ functional connectivity (others) for the working memory task contrasts. We see a diagonal heavy matrix which suggests that there are individual differences in cerebellar-cortical connectivity which is better at predicting the subject’s own task activations as compared to other subjects. b) The distribution of the accuracies made using other subjects’ connectivity (blue) compared to self connectivity (orange).

### Relationship between model coefficients and cortical task activations

Earlier studies (Osher et al., 2019) have shown that the model coefficients are similar to group averaged task activations. We saw similar behavior but with coefficients for predictions in the cerebellum and group averaged task activations in the cortex which were not part of the model dataset. For the contrasts in the Working Memory task, we saw a significant correlation (WM_2BK_BODY: r(398) = 0.62, p<0.0001; WM_2BK_FACE: r(398) = 0.66, p<0.0001; WM_2BK_PLACE: r(398) = 0.68, p<0.0001; WM_2BK_TOOL: r(398) = 0.65, p<0.0001; WM_2BK-0BK: r(398) = 0.70, p<0.0001) between the task activations and ridge regression model coefficients as can be seen in Figure 9 and Figure S3. We saw similar correlations for Social (RANDOM: r(398) = 0.46, p<0.0001; TOM: r(398) = 0.38, p<0.001), Relational (MATCH r(398) = 0.44, p<0.0001), Language (MATH-STORY: r(398) = 0.47, p<0.0001) and Gambling tasks (PUNISH: r(398) = 0.54, p<0.0001; REWARD: r(398) = 0.33, p<0.001) (Figures S4-S6). The activations in the motor regions and coefficients were proportional but the correlations between coefficients and averaged task activations across the whole brain did not come out to be significant. We did not see correlations between the task activations and coefficients for Emotion tasks.

**Figure 9:**
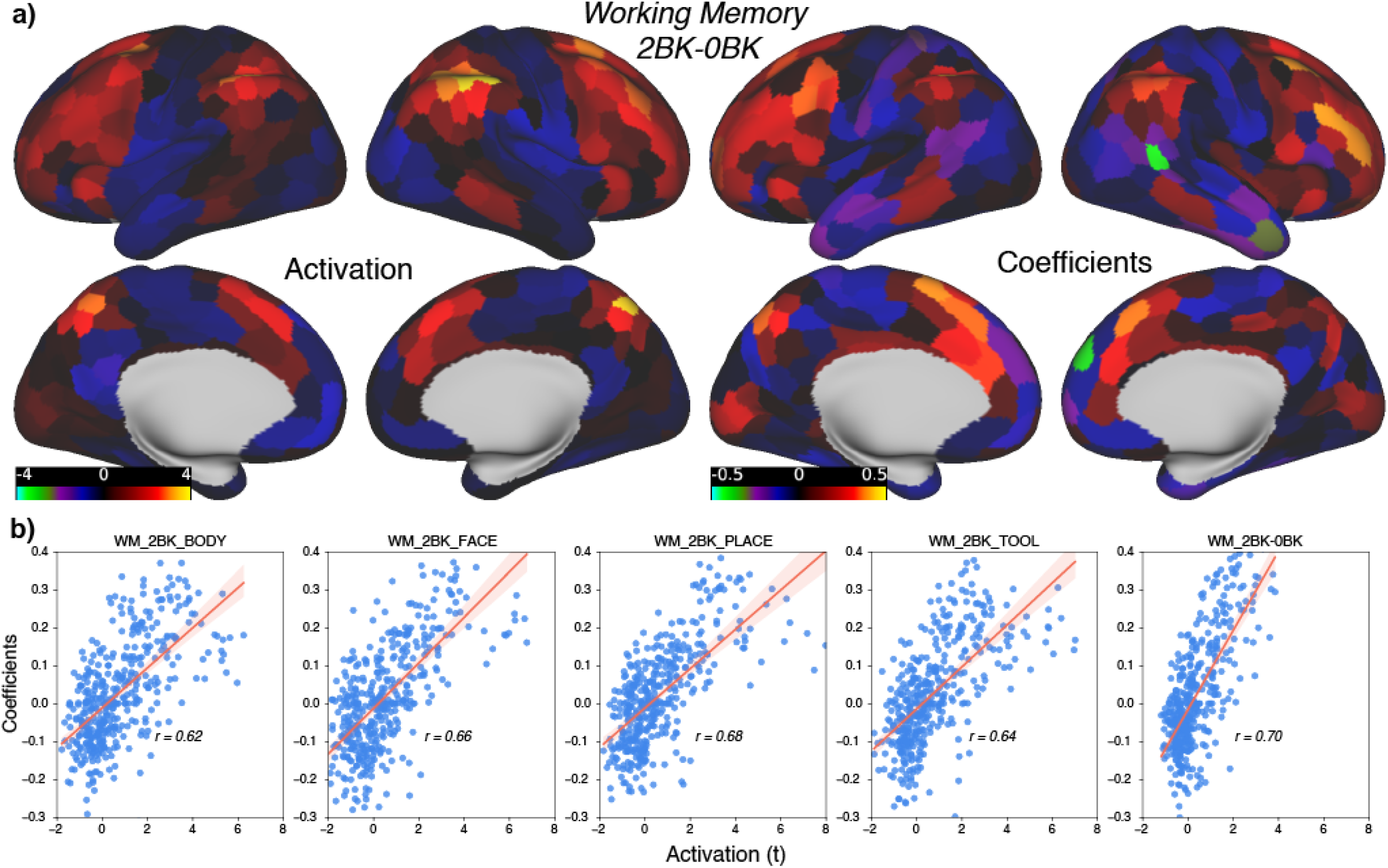
Coefficients and task activations: a) Looking at the distribution of the model coefficients across the cortex for the working memory task (2 back vs 0 back contrast). Group averaged task activations on the left panel and the model coefficients on the right panel. The coefficients have a similar distribution to cortical task activations which were not a part of the training or test model suggesting a link between cerebellar-cortical connectivity and cortical functional areas. b) Scatter plot of model coefficients vs activations (GLM t statistic) for the five contrasts from the working memory task: 2 back body, 2 back faces, 2 back places, 2 back tools and 2 back vs 0 back. A strong association between the model coefficients and the cortical task activations corroborate the link between functional connectivity and function.

## Discussion

There has been an impetus for predictive modeling of the brain activations during cognitive and somatomotor tasks using resting-state functional connectivity (Osher et al., 2019; Tavor et al., 2016; Tobyne et al., 2018). We extended the connectivity fingerprinting based approach for the cerebellum and found that models created using resting-state connectivity with the cerebral cortex perform much better than only group average based predictions across tasks related to various cognitive modalities like working memory, emotion, theory of mind, gambling, relational and motor task. Our work extends the current literature on the association between connectivity between brain regions and the function of brain regions to the cerebellum which for a long time was thought only to respond to motor movements. Recent studies have found cerebellar involvement in attention (Brissenden et al., 2016), working memory (Brissenden et al., 2018, 2021), emotion and other higher cognitive functions (Schmahmann, 1996, 2010, 2019; Stoodley et al., 2012)

We found out that our Connectome Fingerprinting approach predicts task activations significantly better than the group averaged activations in all seven tasks of the HCP dataset. As we limited our search space to task-relevant voxels, some of the contrasts like Language-Math, Language-Story, Gambling_Punish-Reward, Emotion_Faces-Shapes, Social_TOM-Random, Relational_Match-Rel did not have sufficient activation in the cerebellum to be included in the analysis. It could be due to a lack of differences in the activation of these conditions in the cerebellum or across the brain. Our model also did not outperform statistically for the group average model Motor_Tongue-Average, Language_Math-Story and Emotion_Shapes-fix.

Earlier studies have shown the link between the coefficients of CF models trained on cortical search spaces and cortical task activations (Osher et al., 2019). Here, we see that the coefficients of our models based on search spaces defined in the cerebellum were similar to task activations in the cortical regions suggesting a close link between connectivity and function in the cerebellum. Interestingly, we did not see a similar trend for the emotion tasks which could be due to the small number of voxels that were task-relevant in the cerebellum and hence selected in the search space suggesting that the emotion representations in the cerebellum are not strong. The coefficients for motor areas in the cortex were proportional to the cortical motor area activation but not across all the cortical regions. We also saw the specificity of the individual subjects’ functional connectivity in predicting their own activations which other papers have mentioned earlier (Osher et al., 2019; Tavor et al., 2016; Tobyne et al., 2018) suggesting that the cerebellar-cortical connectivity is also individual specific.

Models trained to associate connectivity and function can allow for the prediction of individual-specific functional activations for populations like subjects with ADHD, Alzheimer’s, Dementia or other disorders who can not perform different cognitive tasks in the scanner. It can allow us to predict task-specific regions in clinical populations with brain tumors which can aid in better presurgical planning.

There are a few limitations of the work namely the task design in the HCP protocol. HCP dataset is of very high quality in terms of the acquisition using the latest MR sequences and multi modal registration approaches like the Multi Surface Matching (MSM) (Robinson et al., 2014). But the tasks may not be the best to elicit responses for a particular cognitive modality. The data was also limited by the number of runs per modality. Studies have shown that the more amount of data per subject improves the reliability and the robustness of the results (Braga & Buckner, 2017; Gilmore et al., 2021; Gordon et al., 2017; Somers et al., 2021; Xue et al., 2021). The field has been termed as deep imaging (Gratton & Braga, 2021). We had a large amount of resting-state data per subject but fewer runs of task data per subject which could have resulted in weaker models overall. Our current approach was limited to regularized regression models which perform better than ordinary least squares models (Tobyne et al., 2018) but there might be better performing non-linear models which would require further exploration in future studies.

Our current paper was focused on the tasks in the HCP dataset. We could extend our approach to different datasets which have more focused runs per subject in a specific cognitive modality. We can also perform cross scanner predictions by training on deep imaging subjects and testing them on studies with large cohorts. The combination of models trained on different tasks for the same cognitive modality requires more in-depth research.

## Acknowledgements

Research was supported in part by NSF Grant BCS-1829394 to D.C.S.

## Code/Data Availability

Human Connectome Project dataset is available on db.humanconnectome.org Codes are available on request to the authors.

