## Supplementary figures for "Predicting an Individual’s Cerebellar Activity from Functional Connectivity Fingerprints"

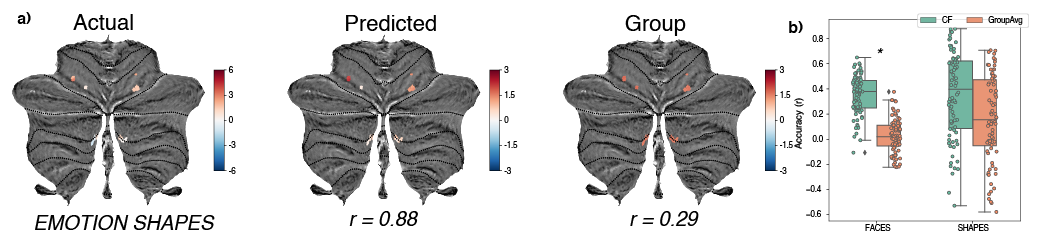

Figure S1: Emotion task Predictions. a) Predictions using the CF model (middle panel) and control Group Average model (right panel) for the emotion task where subjects had to compare either affective faces or shapes. The left panel depicts the actual task activations visualized on the SUIT surface space. b) A very small area of the cerebellum had task-relevant voxels for the emotion task conditions. Prediction accuracies across the test subjects for the two conditions: faces and shapes across the two model types: CF and Group Average. Asterisk depicts statistically significant differences between the prediction accuracies of the two model types.

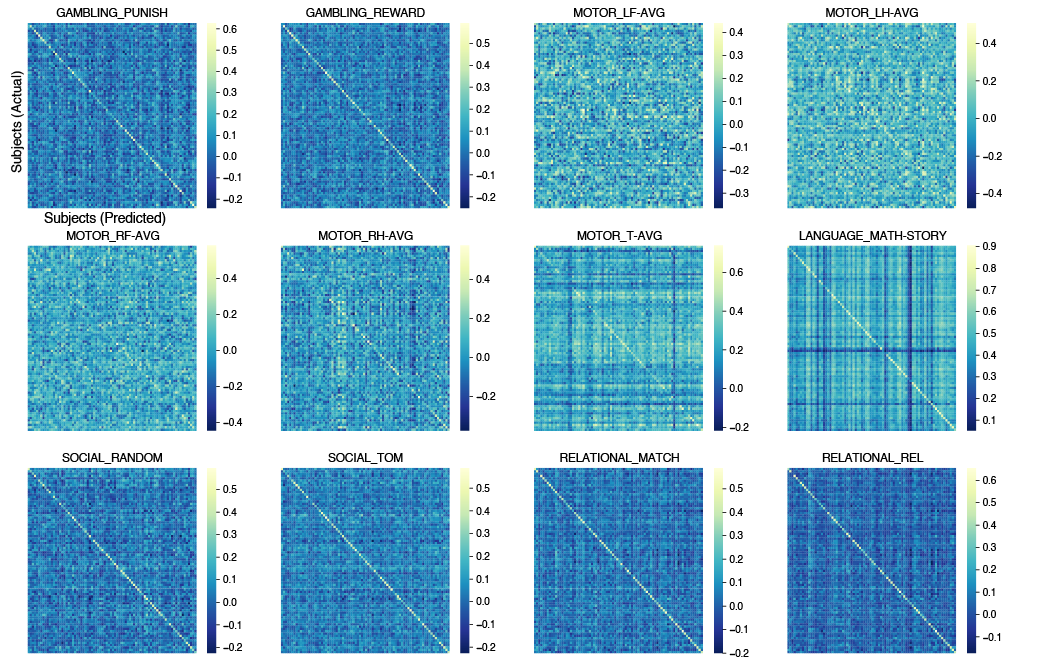

Figure S2:Cross connectivity predictions for the contrasts across the other cognitive tasks. We see that a subject’s own functional connectivity better predicts its task activation as compared to other subjects in gambling, social, relational and language tasks but not so much in the contrasts in the motor task suggesting that cerebellar-cortical connectivity for motor regions is not strongly individual specific.

Figure S3:Coefficients and task activations: Plotting out the surface maps for the coefficients and task activations for the working memory and gambling tasks.

Figure S4:Coefficients and task activations: Plotting out the surface maps for the coefficients and task activations for the motor and language tasks.

Figure S5:Coefficients and task activations: Plotting out the surface maps for the coefficients and task activations for the social, relational and emotion tasks.

Figure S6:Coefficients and task activations: Scatter plots between the model coefficients and task activations for the remaining tasks. Contrasts and tasks with no relationship are not shown here.
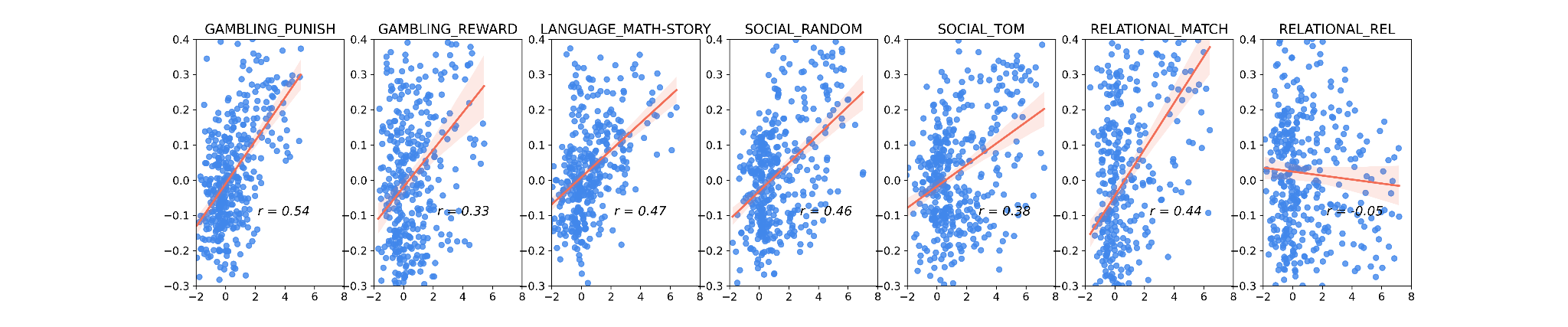
